# Can a Sparse 2^9^ × 2^9^ Pixel Chaos Game Representation Predict Protein Binding Sites using Fine-Tuned State-of-the-Art Deep Learning Semantic Segmentation Models?

**DOI:** 10.1101/2023.08.04.410498

**Authors:** Kevin Dick, James R. Green

## Abstract

No. While our experiments ultimately failed, this work was motivated by the seemingly reasonable hypothesis that encoding protein sequences as a fractal-based image in combination with a binary mask identifying those pixels representative of the protein binding interface could effectively be used to fine-tune a semantic segmentation model. We were wrong.

Despite the shortcomings of this work, a number of insights were drawn, inspiring discussion about how this fractal-based space may be exploited to generate effective protein binding site predictors in the future. Furthermore, these realizations promise to orient complimentary studies leveraging fractal-based representations, whether in the field of bioinformatics, or more broadly within disparate fields leveraging sequence-type data, such as Natural Language Processing.

In a non-traditional way, this work presents the experimental design undertaken and interleaves various insights and limitations. It is the hope of this work that those interested in leveraging fractal-based representations and deep learning architectures as part of their work will benefit from the insights arising from this work.

## Introduction

Many of the biological processes that manifest and maintain life are the result of a complex network of physically interacting proteins. These protein-protein interactions (PPI) play key roles in numerous cellular processes and their elucidation is crucial to both our understanding of cellular processes and disease pathogenesis [1]. Crucial to this task is also the determination of the *region* of a given protein responsible for mediating such interactions: the protein binding site interface. For example, in the case of pathogens, knowledge of these putative binding regions is of particular relevance to disrupt the pathogenic life-cycle. The accurate identification of those amino acid residues responsible for mediating the interactions of a target protein enables the design and development of inhibitory molecules [2], [3], [4].

The development of an effective predictor of binding site interfaces first requires the encoding of a protein sequence into a representation amenable to the use of machine learning models. Numerous encoding methods exist including the position-specific scoring matrix (PSSM) [5], putative relative solvent accessibility [6], evolutionary conservation [6], general numerical descriptors [7], and most recently, the use of protein sequence embedding [8] akin to the word embeddings leveraged in the field of Natural Language Processing (NLP). An increasingly popular method of representing both genomic and proteomic sequences is using the fractal-based Chaos Game Representation (CGR).

### The Chaos Game Representation

The use of CGR to visually depict and represent gene structure was initially proposed by Dr. H. Joel Jeffery in 1990 [9]. Inspired by self-similar fractals, the Chaos Game is an iterated function system (IFS) which incrementally generates an image of the the fractal structure of a sequence of possibly infinite length; that is, a one-dimensional sequence of arbitrary length is encoded into a two dimensional CGR-space with a desirable property that each point in CGR-space encodes both local and global information about the sequence.

The representation for a DNA sequence is created by first defining a four-node square in a plane and assigning to each node one of the four nucleotides from the alphabet Σ = {*A, C, G, T* }. An initial point is placed in the center of the square and then as part of an iterative process, for each nucleotide in the sequence, a line is drawn from the current point towards the node assigned that nucleotide and the current point is moved to the midpoint of that line. This is repeated until the DNA sequence is exhausted. Considering the square plane to be an image of 2^*k*^ × 2^*k*^ black pixels, each pixel visited as part of the Chaos Game is converted to white, generating the CGR image. Each pixel exactly represents a subsequence of length *k* and that pixel is made white if the DNA contains that *k*-mer subsequence and remains black if it does not. This binarized version of the CGR image doesn’t capture the relative frequency of *k*-mer subsequences; an alternative representation that encodes this additional information is known as a *frequency* CGR (FCGR) and is constructed by Min-Max normalizing the frequency count of subsequences where a greyscale value is assigned to the pixel to transform subsequence frequency counts in the range [0, max(*freq*)] ⇒ [0, 1] ⇒ [(0, 0, 0), …, (*i, i, i*), …, (255, 255, 255)] where *I ∈* [0, 255] is an integer encoding the relative subsequence frequency. In prior work, we show that this additional encoded information at times leads to improved predictive performance [10] and can additionally be extened to new domains such as for audio signal representation [11].

In order to represent a protein’s amino acid sequence using the *n* = 4 node representation, we required an effective means of mapping the *n* = 20 amino acid alphabet into the *n* = 4 nucleotide alphabet. To this end, we leveraged the fixed reverse translation mapping proposed by [12] and summarized in Table I (Insight 1).

#### Insight 1

Fixed Reverse Translation Results in Information Loss

The proposed fixed translation mapping proposed by Deschavanne & Tuffery (2008) was selected in such a way as to maintain a balance in base compositions in an effort to maximize differences between amino acid codons [12]. Fundamentally, this mapping may be naïve and fail to necessarily capture biological phenomena or evolutionary features that might manifest at the genomic level yet disappear at the proteomic level. For example, the statistical differences in codon frequencies between organisms are lost in this approach. Moreover, the “third base wobble” may additionally introduce fuzziness within the representation [12]. Since reverse translation is a non-trivial challenge, some loss of information is expected. The work of Deschavanne & Tuffery on protein structural classification reported an approximate accuracy of 84%, rivaling the state-of-the-art at the time.

For a given protein sequence, the reverse translated nucleotide sequence will be three times longer than the original, providing three times as many points in CGR-space when playing the Chaos Game. Consequently, the DNA-based representation will be less sparse than a representation encoding the amino acid sequence itself as part of an *n* = 20 icosagonal representation (Insight 2).

#### Insight 2

Protein Sequence Length ≪ Genome Length

By definition, the sequence length of a given protein is dramatically shorter than an entire proteome, and commensurately shorter than an organism’s genome. By representing individual proteins in this work, the number of characters available within any one sequence are dramatically fewer than the sequences leveraged in prior work where large genomic regions or complete genomes themselves are represented within a single image. For example, previous work by Deschavanne *et al*. successfully characterized and classified species based on whole-genome CGRs [13]. With a sufficiently long nucleotide sequence, CGR is an effective alignment-free means of constructing phylogenetic relationships [14].

Individual protein sequences, even when reverse translated, will only encode a smattering of points in CGR-space as compared to the dense representations leveraged within these prior works. The sparsity of these data can, however, be traded-off with the pixel-level subsequence encoding, discussed next.

**TABLE I.**
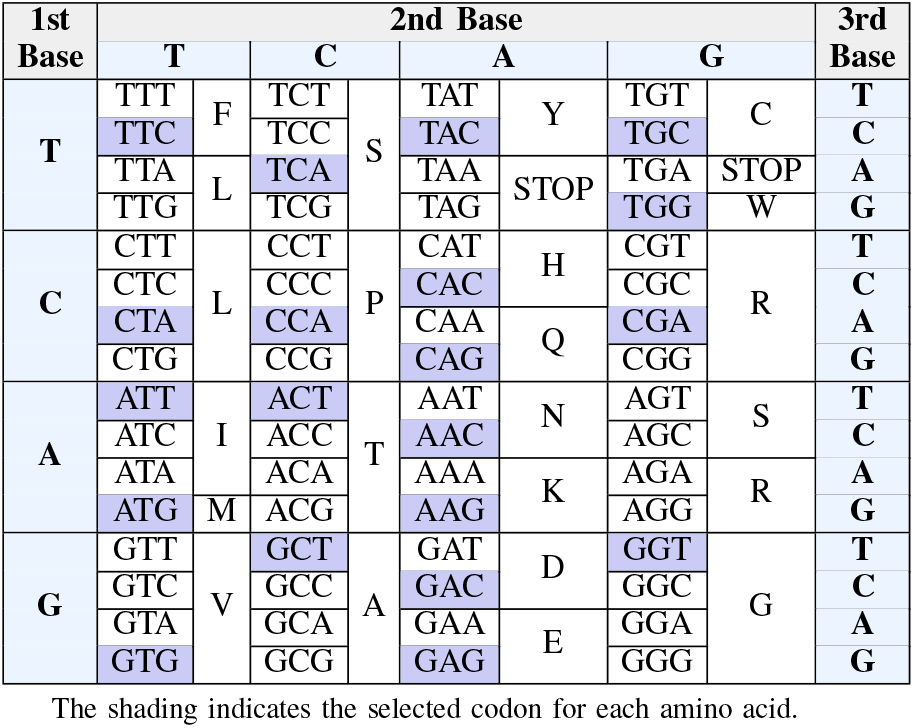
Fixed Reverse Translation Mapping from Dechavanne & Tuffery [12] and Replicated from [10].

Finally, an important consideration when generating CGRs is the selection of an appropriate image size and pixel resolution. As previously described, an image of 2^*k*^ × 2^*k*^ pixels will ensure that each pixel will represent a distinct subsequence of length *k*. This concept is visualized in Fig. 1, where increasing image resolution by orders of two in both directions enables the representation of increasingly long subsequences. While the Chaos Game can be played on non-square images or upon images for which the dimensions are not powers of two, the distinct relationship between a given pixels relative position within an image and the distinct subsequence it encodes is not necessarily conserved. Important to this work is the conservation of this self-similarity in order to conserve a *reverse CGR* mapping of a given pixel back into the sequence domain. By preserving self-similarity, from a given predicted pixel, a quaternary search of *k* steps is needed to recover the predicted subsequence.

**Fig. 1.**
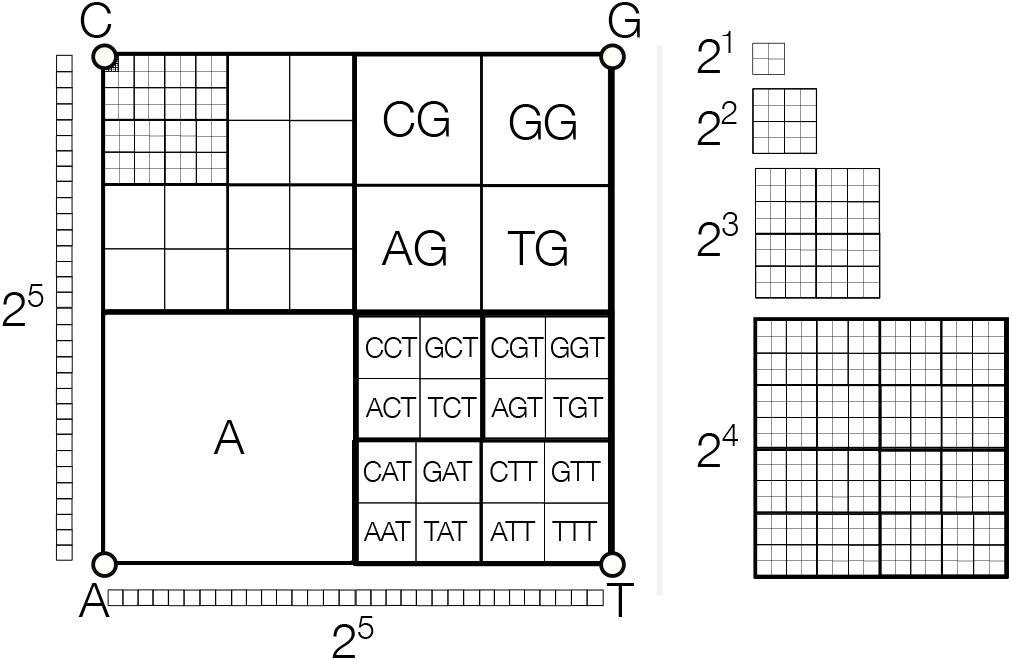
Example Multi-Resolution Image where Distinct Regions Represent Distinct Subsequences. Increasing the pixel resolution by orders of two in both directions conserves self-similarity. Arbitrary image sizes and resolutions are also possible, however, the distinct pixel-position-to-subsequence relationship is not necessarily conserved.

### CGR for Binding Site Interface Prediction

The task of predicting what amino acids contribute to a binding interface requires both the amino acid sequence as well as a ground truth binary mask indicating which of those amino acids indeed participates in the interface (assigned 1) and the remainder which do not (assigned 0). The representation of a sequence in CGR-space is extensible to generating the corresponding binary mask from this ground truth sequence. That is, while generating the amino acid CGR, for each pixel position of the Chaos Game, a separate image encodes the ground truth mask by assigning that pixel to be white if the amino acid participates in the binding interface or remains black (*i*.*e*. unchanged) if it does not. This effectively enables the generation of CGR (image, mask) pairs amenable for use as part of fine-tuning a semantic segmentation model.

Promisingly, each pixel within this CGR-space encodes specific information related to the original protein sequence and, consequently, captures some semantic meaning when combined with the ground truth mask. Represented in this way, these data may be leveraged as part of fine-tuning semantic segmentation task leveraging the state-of-the-art deep CNN architectures.

To that end, we sought to leverage CGR-based representations to fine-tune the state-of-the-art deep CNN models to lerrn the task of predicting the putative binding regions of a given protein.

## Data & Methods

This work used a large and high quality dataset that was originally provided in the work of Zhang *et al*. [6], comprising annotated Uniprot sequences of amino acids, DNA, RNA, and small ligands binding information at the residue level. This dataset was further processed by Li & Ilie as part of [15]. Specifically, only protein-protein binding information was conserved and any sequences sharing more then 40% similarity (measured using CD-HIT) with any of the test sequences were eliminated. Among all proteins of the training dataset, no sequences with similarities greater than 40% were conserved to maximize sequence diversity. Ultimately, the dataset comprises 11,063 protein sequences. For further details on the preparation of this dataset, see [15]. Five independent test sets are additionally considered as part of this work, each denoted as “Dset_72”, “DSet_164”, “DSet_186”, “Dset_355”, and “Dset_448” denoting dataset size. The summary statistics for the length of the protein sequences of each dataset are tabulated in Table II.

For each sample in the training dataset both an amino acid sequence and ground truth binary mask of equivalent length (comprising 1/0s) were provided. This enabled the generation of (image, mask) pairs according to the CGR previously described. Each amino acid sequence was reverse translated according to the mapping proposed by Dechavanne & Tuffery [12]. The binary mask was also extended during the reverse translation. An overview of the (image, mask) encoding process is visualized in Fig. 2 where an amino acid sequence and corresponding mask are reverse translated and respectively extended. The conversion of both into CGR-space required the selection of a resultant image resolution and we choose 2^9^ × 2^9^ pixels to ensure that each pixel represented a nucleotide 9-mer which corresponds to a proteogenic 3-mer (Insight 3).

**Fig. 2.**
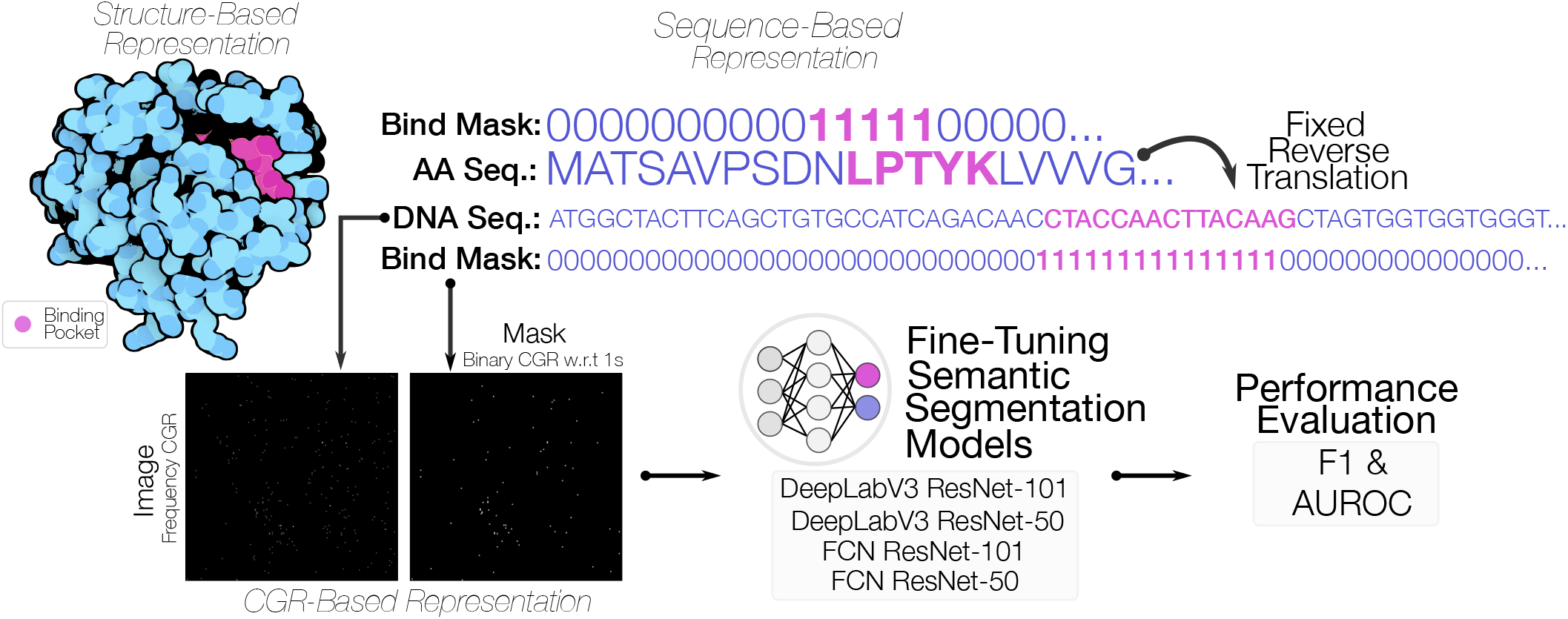
Overview of the Encoding & Prediction Pipeline. For an input amino acid sequence with a binary mask depicting the ground truth amino acids that correspond to a binding interface, the sequence is reverse translated according to Table I and the (F)CGR representation is then generated, producing an FCGR as input image and a binarized mask where each point in the mask corresponds to a 1 in the DNA-encoded binding mask. These (image, mask) pairs are then leveraged to fine-tune four semantic segmentation models originally trained on the ImageNet dataset. Following several training epochs, performance is reported using F1 and AUROC on an independently withheld training dataset.

**TABLE II.** Summary Statistics of the Sequence Lengths for each Dataset.

| Dataset Name | Min. Seq. Length | Max. Seq. Length | Med. Seq. Length | Mean Seq. Length | Std. Dev. Seq. Length |
| --- | --- | --- | --- | --- | --- |
| dset_72 | 45 | 2001 | 175.0 | 252.94 | 256.43 |
| dset_164 | 52 | 870 | 164.5 | 206.37 | 131.60 |
| dset_186 | 40 | 767 | 165.0 | 195.73 | 120.36 |
| dset_355 | 8 | 876 | 256.0 | 272.16 | 135.74 |
| dset_448 | 8 | 876 | 246.5 | 261.72 | 131.70 |
| train_9982 | 8 | 9905 | 306.0 | 427.01 | 489.85 |

### Insight 3

The 2^9^ × 2^9^ Resolution May be Too Sparse

Following from Insight 2, the resulting image from a CGR based on a limited sequence length (single reverse translated protein vs. entire genome) may be too sparse to effectively represent proteogenic information in the minimalist CGR-space. The simple reduction to a consistently self-similar representation of 2^8^ × 2^8^ or 2^7^ × 2^7^ would dramatically increase pixel density.

To corroborate this insight, we plotted the sparsity ratio of each dataset, split into Image and Mask, in Fig. 3. As expected, the masks consistently represent commensurately sparser images given that only a subset of pixels contribute to the actual binding interface. The sparsity ratio is defined as the ratio of the number of zero/black pixels relative to all pixels in the image.

**Fig. 3.**
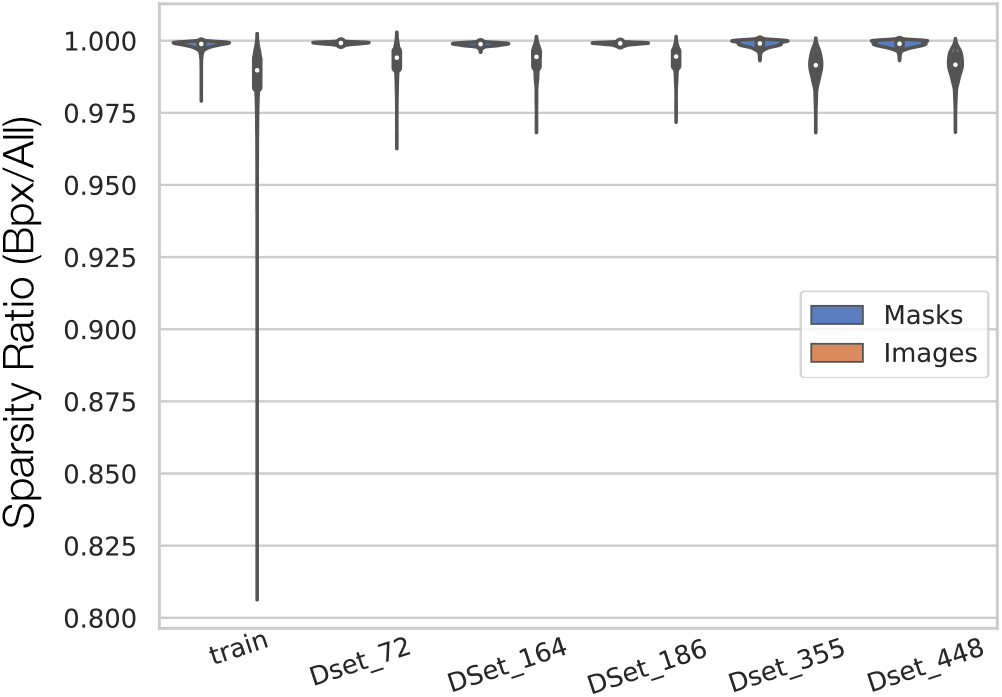
Violin Plots of the Sparsity Ratio across each Dataset. The sparsity ratio is defined as the ratio of zero-based elements compared to encoding pixels. The greater the sparsity ratio, the more sparse a given matrix is.

**Fig. 4.**
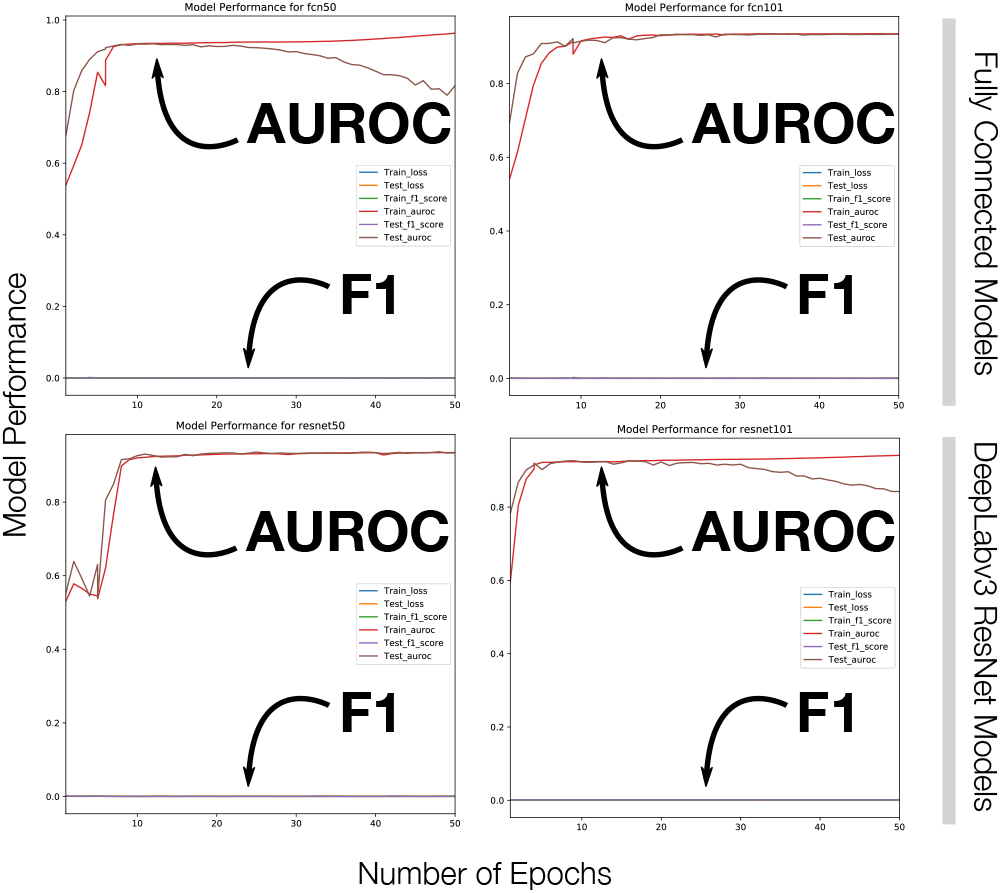
Model Performance Following 100 Epochs of Fine Tuning. Each of the four deep CNN semantic segmentation models failed to capture salient information about the target task. Where both the training and test AUROC climbs as loss drops, the revealing F1 score about zero highlights that the model fails to capture the salient features of the task.

We generated each of the Frequency matrix CGRs (FCGRs) for each protein (capturing the relative frequency of occurrence of *k*-mers) and binary mask CGRs for each reverse translated (AA sequence, binary mask pair) according to Fig. 2. This dataset of (image, mask) pairs was then used as part of a fine-tuning semantic segmentation task to train four models to determine those pixels which contribute to a given protein’s binding interface. Implemented in PyTorch, we considered the ImageNet, pretrained 50-layer and 101-layer Fully Connected Networks (FCN50 & FCN101) as well as Google’s DeepLabv3 ResNet-50 and DeepLabv3 ResNet-101 models.

Each of the four models was fine-tuned over 50 epochs and both the training and testing F1 and area under the receiver-operator characteristic (AUROC) curve metrics were reported.

## Results & Discussion

The task of computationally predicting the subsequence region of interaction for a given protein is of particular importance and novel approaches are of particular interest, should they reveal insights not captured by related perspectives. The consideration of the CGR is one such approach as it simultaneously encoded both local and global information.

We evaluated four state-of-the-art deep CNN models as part of a semantic segmentation task to identify those pixels most likely to contribute to the binding interface and generated models only capable of **predicting all negatives**. That is, over 50 training epochs, our models minimized loss and only learned to classify all pixels as non-interacting (Insight 4).

### Insight 4

The Semantic Meaning of CGR-Space Binding Sites is Fundamentally Different from ImageNet Semantic Meaning

The pretrained models considered in this work were each trained on the ImageNet dataset wherein the proportion of pixels contributing to the semantic meaning of the represented class were both dense and connected. For example, the segmentation of cats within YouTube imagery typically have dense cat-related pixel regions dominant in the view facilitating the transfer learning tasks.

Conversely, this work leveraged a representation that is both sparse and disconnected. This emphasizes the need for sparse- and disjoint-type deep neural architectures amenable to transfer learning tasks of predominantly imbalanced data.

Interestingly, while the training and test loss continually declines over the epochs, the corresponding AUROC typically rises or stabilizes in the near-perfect range. Conversely, the F1 score is consistently hovering about zero. This is a strong indication that our model has learning only to *predict all sites as negatives*; effectively resulting in a null model. This is not entirely unexpected given the extreme sparsity of the images and masks leveraged here (Fig. 3; Insight 5).

### Insight 5

Metric Selection May be Deceiving

While numerous articles tout impressive metrics, they often fail to appreciate the significance of *class imbalance*. That is, when the class of interest is extremely rare, the chosen performance metric should focus on the rare class of interest as opposed to conflating both classes together. In this case, the AUROC of each model is seemingly impressive (near 100%); however, given the class imbalance between actual interface residues and non-interface residues (Fig. 3), the actual metric of interest is the F1 which only marginally hovers above zero. In essence, our models have only learned to classify all residues as negatives!

Whenever a dataset exhibits a modest-to-extreme level of class imbalance (*e*.*g*. 1:10+), in almost no case is a model that predicts only the majority class practical or useful. The *accuracy* and *AUROC* metrics will converge to the value representing the imbalance ratio (IR) within the dataset (1:10 IR ⇒ ∼ 90% Accuracy; 1:1000 IR *⇒ ∼* 99.9% Accuracy). Seemingly impressive; ultimately deceiving.

Typical of class imbalance-presenting problems is the interest in precisely identifying instances of the rare minority class. Thus, metrics capturing our model’s ability to distinguish this class are needed: F1 score and metrics derived from the Precision-Recall curve are ideally suited.

CGR-space differentiates itself from other image-based representations of information in that each pixel represents a specific *k*-mer sequence. As previously mentioned, deep CNNs pretrained on ImageNet are not necessarily able to differentiate between the semantic meaning of pixels in this unique space. Consequently, any patch-level predictions (*i*.*e*. groupings of predicted pixels assigned to the interacting class) will likely contain a considerable number of *invalid* subsequences; that is, *k*-mers which do not actually exist within the original sequence (beyond a reasonable alignment tolerance), further challenging CGR as an effective representation for protein-related tasks. Off-by-one errors in CGR-space are particularly costly as they entirely miss the target site of interaction. Fundamentally, a large-scale dataset of proteins and binding site regions are required to effectively train a model from scratch to learn the physical, chemical, and positional information of a given protein’s putative binding interface (Insight 6).

### Insight 6

More Refined CGRs are Needed

This work emphasizes that semantic segmentation of sparsely distributed pixels cannot be effectively achieved using models which have been trained on images for which the semantic meaning of clusters of pixels are likely to be assigned the same label. In essence, our model learns to predict all pixels as non-interacting which further highlights the inefficacy of certain metrics for evaluation of extremely imbalanced data.

Promisingly, this motivates two subsequent research directions: variation of the image resolution (2 *< k <* 9) as a data augmentation technique and to more densely represent pixels encoding the CGR, and as a compliment, perform end-to-end (from scratch) training of state-of-the-art deep neural architectures using images for which the semantic meaning of pixels are sparsely distributed pixels.

## Conclusion

While this work essentially summarizes negative results, several shortcomings on the use of CGR for general-purpose protein classification tasks were revealed. The CGR has been increasingly adopted as a means to encode features for proteins and may be particularly well-suited for deep neural networks. In addressing many of the open questions and limitations of CGR, it is our hope that this fractal-based representation will be broadly adopted and applied in the field of bioinformatics and more generally within disparate fields reliant on sequence-type data.

Developing effective models for protein amino acid sequences with an | | = 20 alphabet promises further generalization to applications in the field of NLP with |Σ| ≥ 26.

### Proposition

Insight-type Articles to Report Negative Results

While somewhat meta-scientific in nature, the process of reporting on failed work while simultaneously reflecting on the reasons such errors might have manifested by means of interleaved “Insights” may, hopefully, serve as a new model for publishing negative findings. The ability to juxtapose one’s initial assumptions with their realized shortcomings and proposed alternatives for future work is a necessary format in scientific discourse.

From the insights, herein, numerous experiments may be defined to, ultimately, report on the findings originally hoped for this work.

